# FMRI complexity correlates with tau-PET in Late-Onset and Autosomal Dominant Alzheimer’s Disease

**DOI:** 10.1101/2022.06.29.498174

**Authors:** Kay Jann, Julia Boudreau, Daniel Albrecht, Steven Y Cen, Ryan P Cabeen, John M Ringman, Danny JJ Wang, the Alzheimer’s Disease Neuroimaging Initiative

## Abstract

Neurofibrillary tangle pathology detected with tau-PET correlates closely with neuronal injury and cognitive symptoms in Alzheimer’s disease (AD). Complexity of rs-fMRI time-series, measured by entropy values, have recently been reported to decrease with aging, *APOE* ε4 genotype and cognitive decline in AD. Here we hypothesize that the complexity of BOLD signals provides an index for tau-related neuronal injury and cognitive decline in the AD process.

Data were obtained from the Alzheimer’s Disease Neuroimaging Initiative phase 3 (ADNI3) and the Estudio de la Enfermedad de Alzheimer en Jalisciences (EEAJ) study, including cognitively normal elderly controls, persons with late onset AD (LOAD) and early onset autosomal dominant AD (ADAD) patients and their relatives. Our cohort consisted of a sample of 147 subjects from ADNI3 and 41 subjects from EEAJ with T1 structural, tau-PET (tracer: 18F-AV1451) and fMRI scans. Correlations between SUVR tau-PET and multi-scale entropy (MSE) were calculated voxelwise as well as for standard automated anatomical labeling (AAL) atlas regions while accounting for age, gender, and regional gray matter volume. Potential pathways relating MSE to cognitive function mediated through tau-PET were assessed by path analysis.

We found significant negative correlations between low frequency MSE and tau-PET measures in medial temporal lobe, in both ADNI3 and EEAJ cohorts. Furthermore, low frequency MSE showed significant associations with the Clinical Dementia Rating (CDR) scale and the Mini-Mental State Status Exam (MMSE) scores in both ADNI3 and EEAJ cohorts, which were largely mediated through the tau-PET signal.

Correlations of MSE with tau-PET in temporal lobes support our hypothesis that the complexity of rs-fMRI is associated with regional tau protein accumulation. Furthermore, the association of MSE with CDR and MMSE, mediated by tau-PET, in disease relevant areas suggests that a reduction in MSE is indicative of decreased information processing capacity and cognitive decline in AD processes.

## 1. Introduction

With the growing healthcare burden of Alzheimer’s disease (AD) worldwide, biomarkers for pre-symptomatic stages of AD have become increasingly important for the development of preventative interventions ^1^. Based on studies of late-onset AD (LOAD) such as the Alzheimer’s Disease Neuroimaging Initiative (ADNI) study, amyloid-PET imaging of amyloid beta (Aβ) accumulation in the brain is considered an early marker for the preclinical stage of AD (10-15yrs before symptom onset), while tau imaging correlates more closely with neuronal injury and cognitive decline ^2^. Structural and perfusion MRI and FDG-PET have revealed characteristic AD patterns, with volume loss in the hippocampi, and hypoperfusion and hypometabolism in the temporoparietal regions and posterior cingulate cortex (PCC) ^3–5^. These brain areas are also implicated by early and prominent Aβ and tau accumulation. In finding biomarkers for pre-symptomatic AD, fully-penetrant autosomal dominant AD (ADAD) due to *PSEN1, PSEN2,* or *APP* mutations provides a valuable scientific window into the very early stages of AD development ^6–8^. Imaging studies in ADAD such as those by the Dominantly Inherited Alzheimer’s Network (DIAN) show that the overall patterns of Aβ and tau accumulation, atrophy, and hypometabolism/hypoperfusion in ADAD parallel those of LOAD ^9–13^. On the contrary, for tau-PET, a recent DIAN study showed more pronounced accumulation in ADAD subjects as compared to LOAD ^13^. Further highlighting the effect of tau deposition on cognitive decline, both ADNI ^14^ and the Harvard Aging Brain Study ^15^ reported that tau-positive subjects, regardless of their amyloid status, showed more severe cognitive decline than tau-negative subjects. Finally, the level of tau accumulation, but not amyloid, was found to be predictive of the rate of subsequent atrophy. These accumulating findings indicate that tau deposition plays an important role in cognitive decline and neurodegeneration in the progression of AD.

The functional brain architecture of brain networks assessed based on functional MRI has also been associated with the spread of tau accumulation ^16^. Although widely used with promising results in AD ^17, 18^, showing progressively reduced connectivity with increased disease severity, functional connectivity (FC) analysis of rs-fMRI has limited capability to characterize the inherent regional fluctuations of BOLD signal that exhibits dynamic changes within time scales of seconds to minutes (i.e., non-stationary processes) ^19^. There is accumulating evidence that the inherent moment-to-moment variability of BOLD fMRI cannot be treated as mere noise, instead it possesses physiologically meaningful information ^20^ and can provide relevant complementary information to FC ^21^.

In recent years, a variety of complexity metrics derived from the fields of nonlinear statistics and information theory have been developed or adapted to describe the dynamics of physiological systems ^22^, including approximate entropy (ApEn) ^23^, sample entropy (SampEn) ^24^ and multi-scale entropy (MSE) ^25^. Recent studies showed relationships between decreased entropy measures of BOLD fMRI and normal aging ^26–29^, *APOE* ε4 genotype ^30^, as well as with deteriorating cognitive performance associated ADAD ^26^ and mild LOAD ^31^. In particular, MSE analysis at higher time scales or lower temporal frequencies has been shown to be more sensitive in detecting aging effects and AD progression by filtering out random fluctuations at higher temporal frequencies.

Based on the emerging associations between cognitive decline and tau-accumulation as well as reduction of entropy, we formulated the working hypothesis that the complexity of BOLD signals at higher time scales (i.e. slower frequencies) provides an index of the information processing capacity of regional neuron populations, and is therefore sensitive to tau-related neuronal injury and cognitive decline in the processes of AD.

## 2. Methods

### 2.1 Subjects

#### 2.1.1 Late Onset AD (LOAD) cohort: ADNI3

Data used in this study were obtained from the ADNI database (adni.loni.usc.edu). The ADNI was launched in 2003 as a public-private partnership, led by Principal Investigator M. W. Weiner, MD. The primary goal of ADNI has been to test whether serial MRI, PET, other biological markers, and clinical and neuropsychological assessment can be combined to measure the progression of MCI and early AD. In our study we evaluated the relationship of tau-PET and complexity of rs-fMRI and thus we identified ADNI participants from phase 3 (ADNI3) that had a tau-PET scan (tracer: 18F-AV1451), a rs-fMRI scan and a T1w structural scan at baseline. Resting state fMRI (rs-fMRI) was acquired with the ADNI basic protocol: gradient-echo echo-planar imaging (GE-EPI) with isotropic voxel size of 3.4×3.4×3.4mm^3^, TR/TE=3000/30ms, acquisition time ~10min with 197 volumes. We identified a cohort of 147 subjects, with mean(+/-std) age of 72.4(7.7) years and gender distribution of 82F/65M. Cognitive assessment in this ANDI3 cohort classified 95 as cognitive normal (CN), 15 with Early Mild Cognitive Impairment (EMCI), 7 with Late Mild Cognitive Impairment (LMCI), 23 with MCI and 7 with mild AD. Details of clinical diagnosis in ADNI have been previously described (^32, 33^ for general methods, see http://adni.loni.usc.edu/adni-3/). Briefly, CN participants had a Mini-Mental State Examination (MMSE) score of >26 and a Clinical Dementia Rating (CDR) of 0. Participants with probable AD had an MMSE score <24, and a CDR of 0.5 or 1.0. MCI, EMCI, and LMCI were pooled into one MCI group (N=45). Detailed demographic and clinical information for the three ADNI3 groups can be found in Table1.

**Table 1:** Demographic of ADNI and EEAJ cohorts. (MMSE Mini Mental State Exam; CDR Cognitive Dementia Rating; CN Cognitive Normal; MCI Mild Cognitive Impairment; AD Alzheimer’s Disease)

| ADNI | CN | MCI | AD | statistic | significance | effect |
| --- | --- | --- | --- | --- | --- | --- |
| **N** | **95** | **45** | **7** |  |  |  |
| **age (mean/std)** | 72.9 (8.0) | 72.2 (6.8) | 67.3 (8.0) | F=1.8 | p=0.168 | n.s |
| **gender (m/f)** | 57/38 | 23/22 | 2/5 | Chi<sup>2</sup>=3.2 | p=0.203 | n.s |
| **MMSE** | 28.9 (1.36) | 28.2 (2.07) | 22.4 (2.99) | F=47.7 | p<1e-15 | CN>MCI>AD |
| **CDR** | 0.03 (0.11) | 0.38 (0.27) | 0.79 (0.39) | F=89.3 | p<1e-25 | CN<MCI<AD |
| EAAJ | CN | MCI | AD | statistic | significance | effect |
| **N** | **20** | **13** | **8** |  |  |  |
| **age (mean/std)** | 30.6 (6.7) | 41.8 (9.3) | 46.3 (11.4) | F=12.5 | p<0.0001 | CN<(MCI=AD) |
| **gender (m/f)** | 10/10 | 8/5 | 4/4 | Chi<sup>2</sup>=0.48 | p=0.788 | n.s |
| **MMSE** | 28.8 (1.44) | 19.8 (6.27) | 12.3 (5.47) | F=43.0 | p<1e-09 | CN>MCI>AD |
| **CDR** | 0.0 (0.00) | 0.81 (0.25) | 1.88 (0.64) | F=108.2 | p<1e-15 | CN<MCI<AD |

#### 2.1.2 Autosomal Dominant AD (ADAD) cohort: EEAJ

We also included a second cohort of participants from the Estudio de la Enfermedad de Alzheimer en Jalisciences (EEAJ, PI John M Ringman), a study that is focused on understanding Alzheimer’s in persons of Mexican Mestizo origin that are at-risk for fully-penetrant ADAD (due to either the A431E, or I180F *PSEN1* or V717I *APP* mutations). This dataset included 20 CN, 13 MCI and 8 AD subjects with mean(+/-std) age of 37.2(10.7) years and gender distribution of 19F/22M. Cognitive status was determined by a comprehensive clinical assessment including formal neuropsychological testing according to the UDS3 of the NIA-funded Alzheimer’s Disease Centers ^34^ including MMSE and CDR. Diagnoses were rendered according to the UDS protocol ^34, 35^ adjudicated by author JMR. These study participants had a T1w structural scan, a tau-PET scan (tracer: 18F-AV1451) scan and two rs-fMRI sessions with opposite phase encoding (AP/PA) based on the Human Connectome Project protocol using a multiband (MB) GE-EPI sequence with isotropic voxel size = 2×2×2mm^3^, TR/TE=720/33ms, MB factor=8, and acquisition time of 5min with 420 volumes. Detailed demographic information for the three EEAJ groups can be found in Table1.

ADNI3 or EEAJ studies are in accordance with the Declaration of Helsinki and approved by the respective institutional review boards.

Data availability is provided through the ADNI and EEAJ steering committees and data use agreements.

### 2.2 Data preprocessing

#### 2.2.1 Resting-state fMRI data

FMRI data were motion-corrected, normalized to Montreal Neurological Institute (MNI) canonical space and smoothed with an 8mm Gaussian kernel. Physiological and motion related signal fluctuations were regressed out based on eroded WM and CSF masks, generated from probabilistic tissue segmentation masks of T1w-images, and 12 motion parameters (x,y,z translation and rotation plus first derivatives), respectively. MSE was computed using the in-house developed LOFT Complexity Toolbox (github.com/kayjann/complexity). Different time scales were calculated by coarse-grained averaging of the original BOLD time-series. Temporal frequency was calculated by 1/(scale*TR), thus low-scales represent higher frequency complexity (scale 1 is the original temporal resolution) while higher scales capture complexity of low frequency signal fluctuations. In total we calculated 6 scales in each cohort with pattern matching length m = 2 and pattern matching threshold r = 0.50. The choice of these parameters was in the reported range of values for fMRI data in literature ^36, 37^ and based on previous work in our lab ^27, 31, 38^. As previous research in AD demonstrated that AD, MCI, and CN show the largest differences in high-scale entropy (low-frequency), we limited further analysis to the highest scale in each cohort. For the EEAJ cohort MSE was calculated for AP and PA scans separately and then averaged into a single map.

#### 2.2.1 Tau-PET data

Tau-PET data were normalized into MNI space and smoothed with an 8mm Gaussian kernel. Cerebellar segmentation was performed with SUIT (http://www.diedrichsenlab.org/imaging/suit.htm), and dorsal regions were removed from the cerebellar ROI ^39^. Average PET signals were extracted for reference regions in inferior cerebellar gray matter in native PET space. Parametric Standardized Uptake Value Ratio (SUVR) maps were created by dividing the PET signal in each voxel by the average signal in the cerebellar reference region.

### 2.3 Statistical Analysis

For the ADNI and EEAJ cohort, we first calculated the average MSE and tau-PET SUVR maps for each group: CN, MCI, and AD. This provided a visual representation of the spatial pattern of changes in each metric with progressive disease severity. We expected to see decreases in rs-fMRI MSE along with increases in tau-PET SUVR values in brain areas known to be involved in disease progression such as the medial and inferior temporal lobe, PCC, and potentially parietal and frontal cortex, which will be a confirmation of data consistency with the current literature

#### 2.3.1 Association between rs-fMRI MSE and tau-PET SUVR

To assess the relationship between rs-fMRI MSE and tau-PET SUVR we performed voxel-wise partial correlation analysis including age, gender, and regional gray matter volume as covariates. Regional gray matter volume was calculated based on individually segmented T1w-images using SPM12’s Dartel algorithm. This analysis was performed for the ADNI and EEAJ cohorts independently. Statistical significance was set at p<0.05 and multiple comparison correction was applied using a cluster-size threshold estimation in alpha-sim (www.nitrc.org/projects/rest/) using 1000 permutations. In addition to the voxel-wise analysis, we repeated the correlation analysis for larger regions of interest (ROIs) based on parcellation of the cortex into the AAL atlas ROIs. This approach benefited from improved signal-to-noise ratio (SNR) for both MSE and SUVR. Results for both voxel-wise and ROI-based analyses were projected onto cortical surfaces for visualization using the Quantitative Imaging Toolkit (QIT, https://cabeen.io/qitwiki,^40^).

#### 2.3.2 Association between rs-fMRI MSE and cognitive scores

Based on the findings in the voxel-wise analysis we identified three clusters with significant voxel-wise correlations that were located in disease relevant cortical areas. From each of these clusters we extracted the average MSE values to calculate the association with CDR and MMSE scores, respectively. This analysis was based on partial correlation analysis including age, gender, and average regional gray matter volume as covariates and displayed using scatter plots with adjusted values. Residual plots were used to assess the model integrity. Correlation analyses were performed using the Statistics and Machine Learning Toolbox in MatLab.

Furthermore, we performed mediation analyses in these same clusters to evaluate the contribution of tau-PET on the relation between MSE and MMSE or CDR respectively. Mediation effects ^41^ were tested in SAS software and statistical significance was tested by 95% confidence-interval estimated with 1000 bootstrapping samples.

## 3. Results

### 3.1 Entropy and tau-PET

Figure1 displays the average maps for MSE at the highest scale as well as tau-PET SUVR for each subgroup of the ADNI and EEAJ cohort, respectively. It can be seen that in both cohorts MSE is higher in the NC compared to MCI and then AD groups (Fig1 A & C). Notably, prominent MSE reduction can be observed in temporal lobes of MCI and AD groups compared to NC. Similarly, a gradual reduction of MSE with increasing disease severity can be observed in parietal and frontal cortices. In contrast, tau-SUVR manifests an increase from NC, MCI to AD, notably in inferior temporal areas and PCC along with slower increases in the frontal cortex (Fig1 B & C). Furthermore, tau-PET deposition appears more severe in the ADAD-EEAJ cohort compared to the ADNI cohort. These qualitative findings demonstrate an overall inverse association between MSE and tau-PET, which will be elaborated in the correlation analysis below.

**Figure 1:**
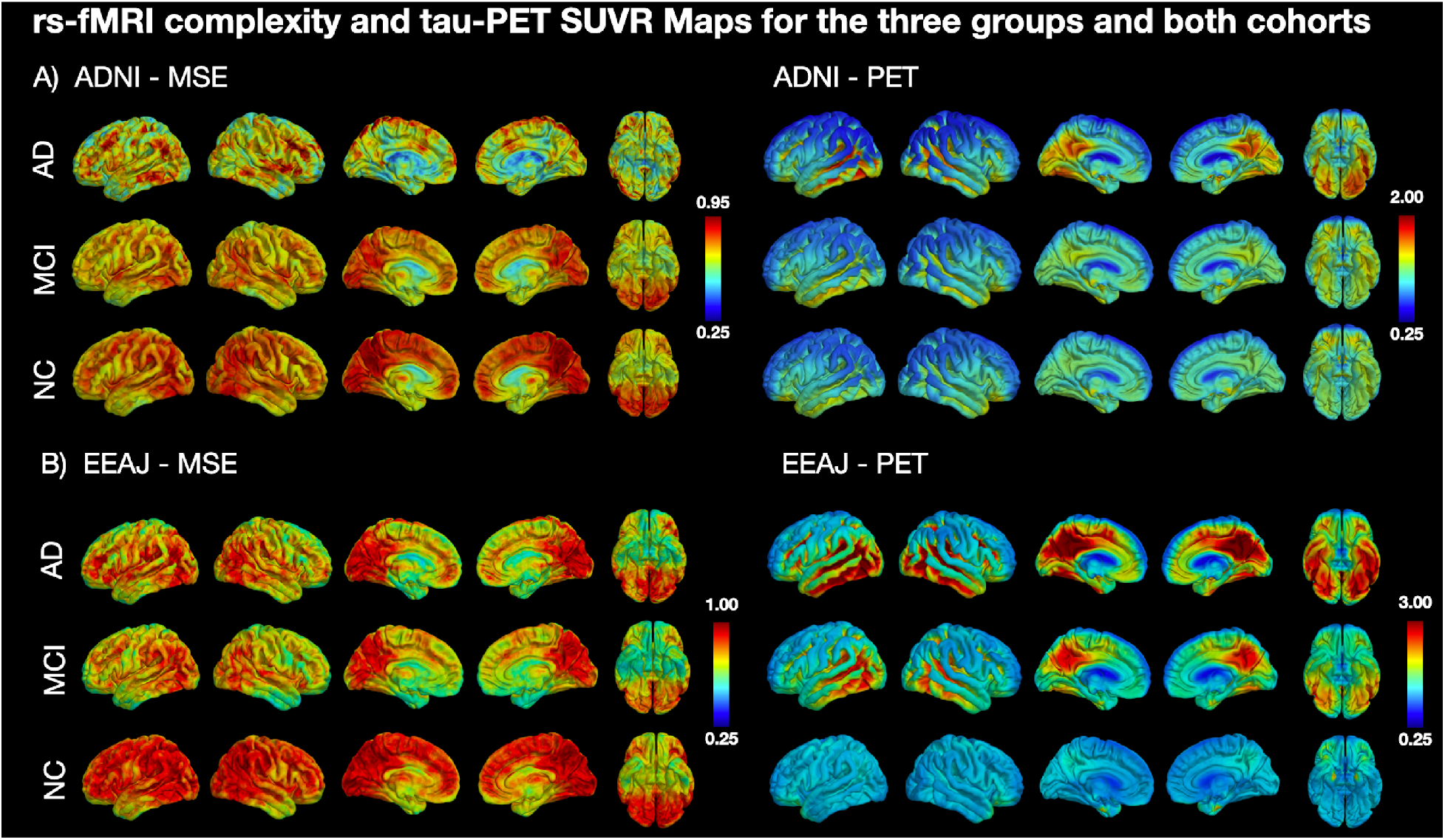
Mean maps for rs-fMRI complexity and tau-PET SUVR in subgroups of both cohorts. (CN Cognitive Normal; MCI Mild Cognitive Impairment; AD Alzheimer’s Disease; MSE Multi-Scale Entropy).

### 3.2 Association between Entropy and tau-PET

To estimate the association between MSE and tau-PET SUVR, we calculated Spearman correlations between the two measures, including age, gender, and regional gray matter volume as covariates. Additionally, this analysis was performed on a voxel-wise level as well as for larger ROIs based on the AAL atlas parcellation. The voxel-wise analysis revealed a number of significant negative correlations between MSE and tau-PET in the ADNI cohort. Notably, there were clusters in lateral and inferior temporal lobe, superior parietal lobe, and lateral frontal lobe (Fig2A). For the EEAJ cohort, we found significant negative correlations in bilateral parahippocampal gyri, lateral temporal lobe, anterior cingulate cortex (ACC), and lateral frontal lobe (Fig2B). There were no significant positive correlations in either cohort. The ROI-based analysis revealed significant negative correlations in left hippocampal gyrus, bilateral lingual gyrus, and occipital lobe for the ADNI cohort (Fig2A). The EEAJ cohort also showed strong significant negative correlations in bilateral parahippocampal, fusiform gyrus, as well as ACC, precuneus and right lateral frontal lobe (Fig2B). Again, there were no areas with positive correlation in either cohort. We further extracted the values from the clusters with significant results in the voxel-wise analysis and display the relationship between MSE and tau-PET in Figure 3. Overall, these results confirmed negative relationships between rs-fMRI MSE and tau-PET deposition in areas associated with AD progression. In the ADNI cohort we observed statistically significant negative associations in the precuneus cluster (r=-0.25 p=0.003) and two clusters in the inferior temporal lobe, left inferior temporal gyrus (ITG) (r=-0.31 p=0.0002) and right inferior temporal gyrus (ITG) (r=-0.35 p=0.00002). In the EEAJ cohort, all three ROIs also exhibited strong negative significant correlations: ACC (r=-0.45 p=0.005), left parahippocampus (r=-0.50 p=0.002), and right parahippocampus (r=-0.53 p=0.0006).

**Figure 2:**
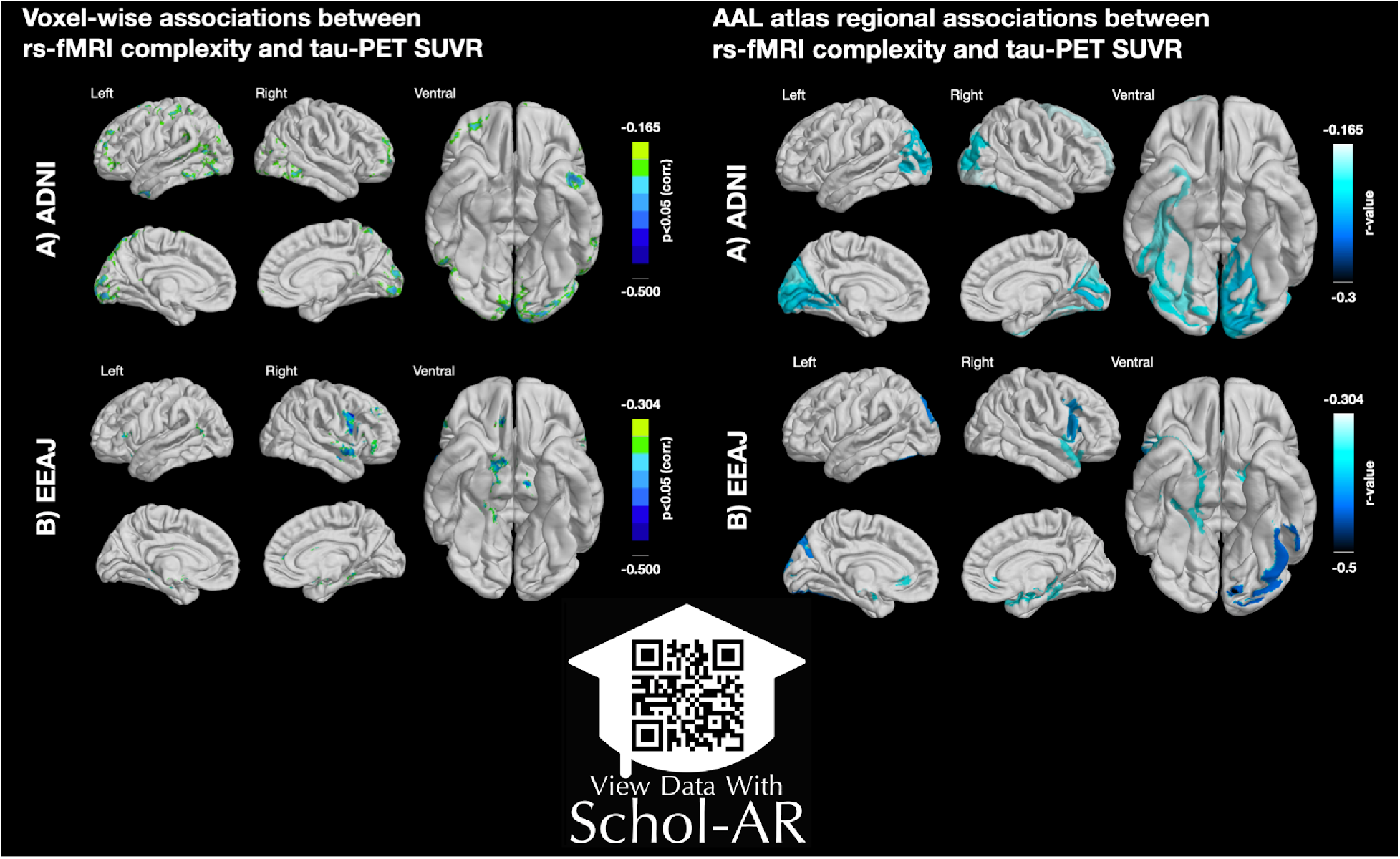
Left panel: Voxel-wise associations between rs-fMRI complexity and tau-PET SUVR. Right panel: AAL atlas regional associations between rs-fMRI complexity and tau-PET SUVR. (AAL automated anatomical labeling atlas). ScholAR QR code will enable augmented reality 3D visualization of the results displayed in the left panel using the Schol-AR app or opening manuscript pdf in www.schol-ar.io/reader.

**Figure 3:**
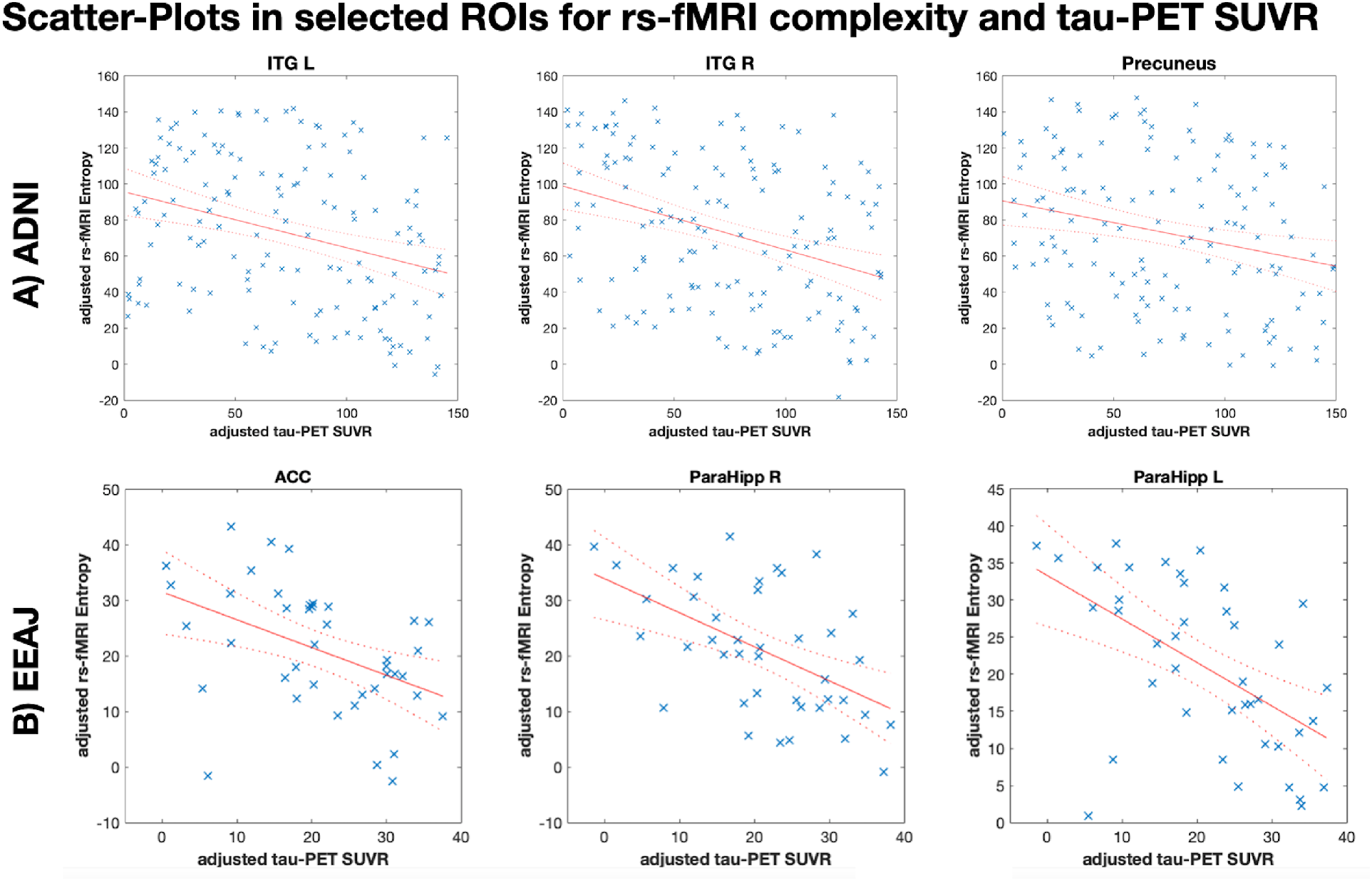
Scatter-plots in three selected regions of interest (ROIs) displaying the relation between rs-fMRI complexity and tau-PET SUVR. (ACC: anterior cingulate cortex); ITG R/L inferior temporal gyrus left/right; ParaHipp L/R parahippocampal gurys left/right)

### 3.3 Correlation and mediation analysis for cognitive decline

To confirm the link between alterations in MSE and cognitive decline, we performed partial Spearman correlation analyses for CDR and MMSE against MSE, respectively, accounting for age, gender, and regional GM as covariates (Figure 4). We used the same clusters from the voxelwise analysis that showed significant associations between MSE and tau-PET in section 3.2. For CDR we found statistically significant negative correlations for all ROIs in ADNI (Precuneus r=−0.37 p<0.0001, left ITG r=-0.18 p=0.029, right ITG r=-0.27 p=0.001) and for left parahippocampus in EEAJ (FIG4A). ACC (r=-0.35 p=0.025) and right parahippocampus (r=-0.34 p=0.031) did show negative correlation between MSE and CDR in EEAJ but did not reach significance. For MMSE (Fig4B), we observed weak positive correlations in the ADNI cohort only reaching significance in the precuneus cluster (r=0.25 p=0.002) but not in the two inferior temporal lobe areas (left ITG r=0.03 p=0.659, right ITG r=0.14 p=0.094). In contrast, the EEAJ cohort displayed highly significant positive correlations between MMSE and MSE in ACC (r=0.52 p=0.001), and bilateral parahippocampal clusters (left r=0.55 p=0.0004, right r=0.39 p=0.016). These findings indicate that with progressing cognitive decline there is a decreasing complexity of rs-fMRI BOLD signal.

**Figure 4:**
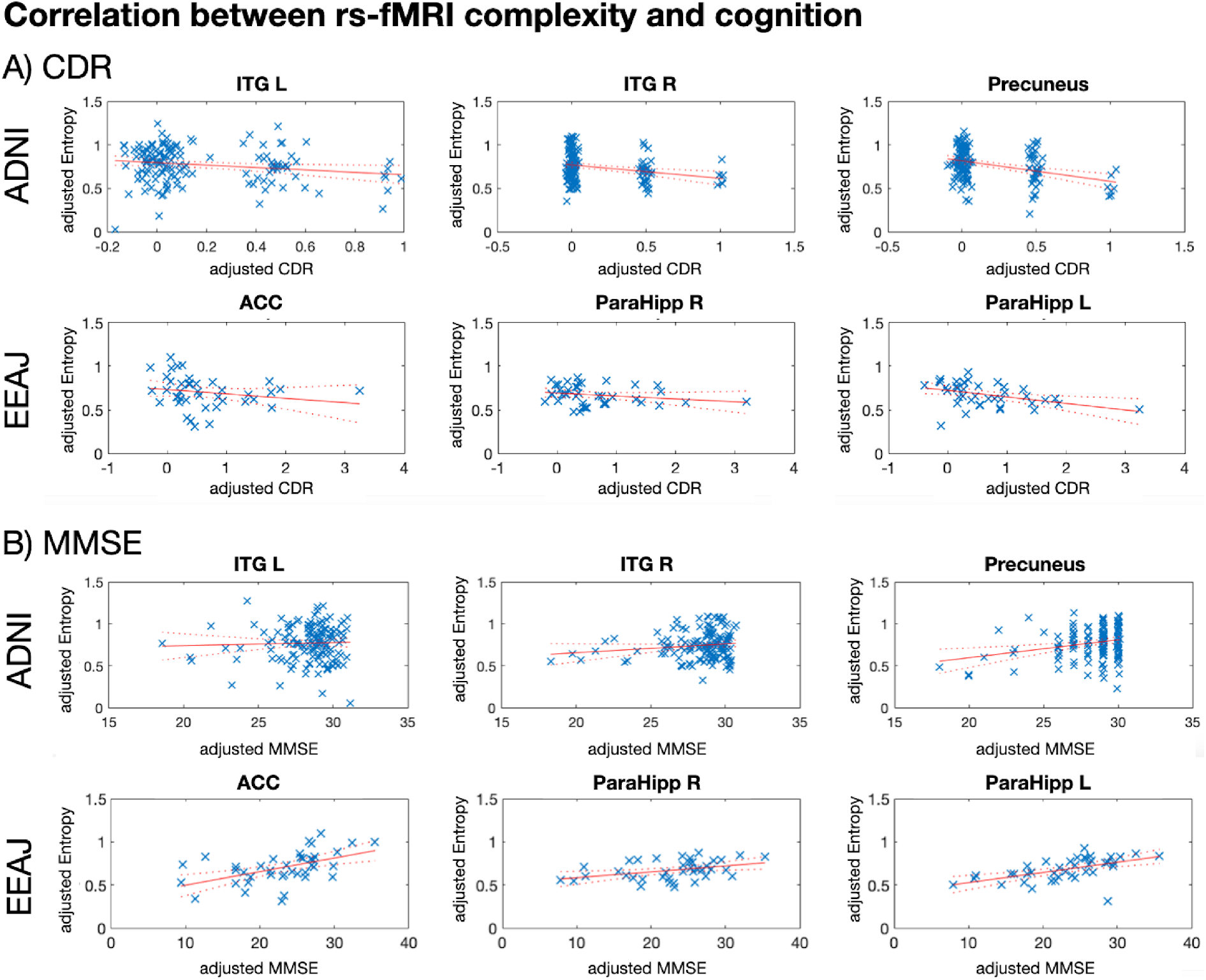
Correlation between rs-fMRI complexity and cognition. ((MMSE Mini Mental State Exam; CDR Cognitive Dementia Rating; ACC: anterior cingulate cortex); ITG R/L inferior temporal gyrus left/right; ParaHipp L/R parahippocampal gurys left/right)

The mediation analysis for ADNI data revealed statistically significant indirect (mediation) effects of tau-PET (0.03 95% confidence interval (CI) (0, 0.06)) on the total association between MSE and log-transformed MMSE (0.13 95% CI (0.05, 0.2)) in the precuneus ROI. The two inferior temporal ROIs did not show statistically significant mediation effects for MMSE (Left: total 0.01 95% CI (−0.05, 0.08), indirect 0.02 95% CI (−0.02, 0.05), Right: total 0.08 95% CI (−0.01, 0.17), indirect 0.04 95% CI (−0.01, 0.09)). Similarly, precuneus showed statistically significant mediation effect for CDR of −0.41 95% CI (−0.99, −0.11) from the total effect of −2.78 95% CI (−4.04, −1.52), while the two temporal ROI did show a statistically significant total effect but not the mediation effect (Left: total −1.17 95% CI (−2.28, −0.05), indirect −0.18 95% CI (−0.74, 0.15), Right: −2.05 95% CI (−3.54, −0.56), indirect −0.18 95% CI (−0.75, 0.38)).

In the EEAJ cohort, mediation analysis revealed statistically significant indirect (mediation) effects of tau-PET on the total association between MSE and MMSE (log-transformed) in both parahippocampi: Left: total 1.57 95%CI (0.75, 2.39), indirect 0.83 (0.15, 1.81) and Right: total 1.32 (0.19, 2.44), indirect 1.43 (0.45, 2.61), as well as in ACC: total 0.92 95% CI (0.3, 1.54), indirect 0.52 95% CI (0.11, 1.1). Furthermore, CDR showed statistically significant associations with MSE with mediation by tau-PET in left parahippocampus total −2.17 95% CI (−3.84, −0.51), indirect −1.45 95% CI (−2.64, −0.42). For right parahippocampus and ACC the total effect was not significant but mediation (indirect effect) showed significant effects: right parahipp: total −1.43 95% CI (−3.6, 0.74), indirect −2.29 95% CI (−3.81, −0.86), and ACC total −0.78 95% CI (−2.09, 0.52), indirect −1.09 95% CI (−2.23, −0.36).

Figure 5 shows a conceptual model of mediation effect and summary of mediation analysis.

**Figure 5:**
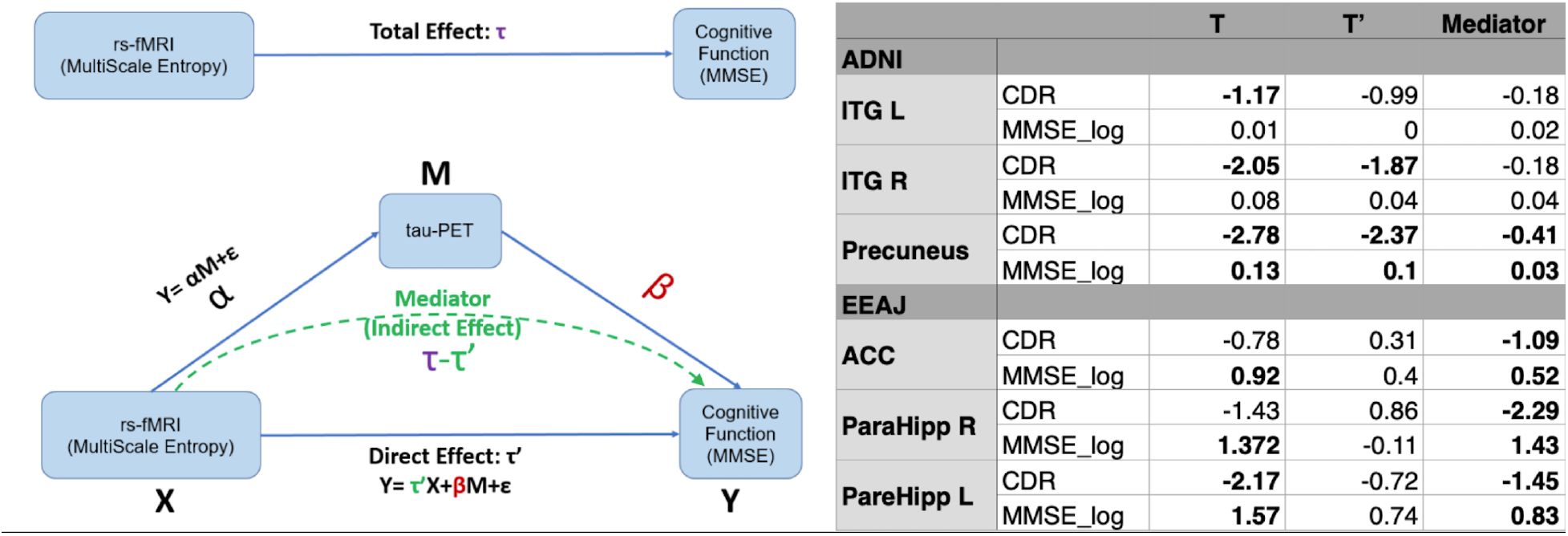
Conceptual visualization of mediation effect modeling and summary table of mediation analysis. ((MMSE Mini Mental State Exam; CDR Cognitive Dementia Rating; ACC: anterior cingulate cortex); ITG R/L inferior temporal gyrus left/right; ParaHipp L/R parahippocampal gurys left/right)

## 4. Discussion

In this study we tested a hypothesis on the relationship between tau deposition and complexity of rs-fMRI BOLD signal fluctuations. This hypothesis was motivated by growing literature indicating that the accumulation of tau protein in specific brain areas is related to cognitive decline in AD ^13, 14, 42^ and literature suggesting that complexity of rs-fMRI signal decreases is associated with aging, *APOE* genotype, and cognitive decline ^27, 30, 37^. These data prompted our hypothesis that areas manifesting increases in tau deposition measured by tau-PET should exhibit reduced complexity of rs-fMRI measured by MSE. We focused on the entropy of low-frequency fluctuations which have been reported to be most sensitive to cognitive decline by filtering out noise-like high frequency fluctuations. Our main findings in the two independent cohorts revealed an exclusively negative correlation between tau-PET and MSE and decreases in MSE with more severe cognitive impairment in AD-associated brain areas, thus confirming our overall hypothesis.

For the ADNI and EEAJ cohorts there was a visible increase in tau-PET SUVR in temporal, parietal, and frontal areas consistent with the literature and AD disease progression models. For MSE, we report high overall complexity in the normal controls and continuously reduced complexity in the MCI to AD groups. This reduction was especially pronounced in the temporal lobe and areas of the frontal lobe. Correlational analysis between tau-PET and MSE at the voxel level revealed sparse but significant negative correlations in the temporal cortex in both cohorts and additionally in the anterior cingulate cortex in the EEAJ cohort.

Temporal and parietal regions have been most consistently reported to show entropy reductions in ADAD and LOAD populations, including middle and superior temporal gyrus, supramarginal gyrus, precuneus, and PCC. In our latest study, using an independent sample of NC and mild AD, we also found that the earliest reduction of MSE in the hippocampus is associated with reduced connectivity within the DMN and cognitive decline ^31^. The spatial pattern of entropy reduction and correlation with tau-PET in the ROI analyses matches the characteristic pattern of temporoparietal hypometabolism in AD detected by FDG-PET as well as higher tau protein accumulation in the temporal cortices of MCI/AD subjects ^13, 43, 44^.

We found a stronger association between MSE and tau-PET in the ADAD EEAJ cohort, while changes in fMRI complexity with disease progression were similar but weaker in the LOAD ADNI cohort. In particular, ADAD can be viewed as a “pure” form of AD without the comorbidities of aging and declining vascular health present in LOAD ^19, 26, 45^. The ADNI cohort had by design much lower severity of cognitive impairment and participants were of advanced age such that additional comorbid pathology independent from tauopathy could have effects on fMRI complexity estimates. Thus, the weaker association between fMRI complexity and tau deposition could have been expected. On the other hand, the autosomal dominant EEAJ cohort was younger in age and neurovascular factors less prevalent, and we observed a much closer association between complexity and tau deposition in AD associated brain areas, specifically the parahippocampal gyrus in the voxel wise comparison, but also in the ROI based results. Another possible contribution to the stronger association observed in the ADAD population is the wider range of AD disease severity in this group. Overall, the observed differences of weaker association between fMRI complexity and tau-PET in LOAD compared to ADAD populations could indicate that vascular or other factors associated with aging may contribute substantially to cognitive decline in LOAD in combination with tau burden ^5^.

To confirm the association of fMRI complexity and tau-PET to global cognitive decline in AD, we performed correlation and mediation analyses in select ROIs. We found that MSE is positively correlated with fMRI complexity and negatively correlated with CDR. Notably, these associations were most striking in bilateral parahippocampus in the EEAJ cohort, corroborating our speculations on reduced bias of comorbid pathology in the ADAD as compared to LOAD. Furthermore, our findings are consistent with two recent studies that used rs-fMRI data from the ADNI study, employing MSE and permutation entropy (PE) analyses and reporting progressive reduction of entropy from normal controls (NC) and early MCI (EMCI), to late MCI (LMCI) and AD groups, with significant associations between complexity measures of rs-fMRI and cognitive decline in MCI/AD subjects ^37, 46^. Together, our findings and previous studies strongly support the notion that with increasing cognitive decline, there is a reduction of fMRI complexity. Mediation analysis provided further insight into this relationship demonstrating that the association between MSE and cognition is mediated by underlying tau pathology. Compared to previous studies that only looked at the association of complexity to cognitive decline, we provide the first evidence that tau accumulation is the mediating factor between these metrics. Accordingly, tau related neurotoxic processes alter neuronal and neurovascular function, which ultimately lead to neurodegeneration. However, it has been reported that tau deposition and changes in neuronal activity and blood flow precede neurodegeneration and patients display cognitive impairment before detectable structural changes in MR. We propose that fMRI complexity provides a new means to assess neurofibrillary tangle induced dysfunction. While neurofibrillary tangle pathology can be seen as the underlying physiological mechanism causing changes in neuronal function and ultimately cognitive decline, changes in fMRI complexity represent an associated functional phenomenon in the AD disease progression.

There are several limitations to this study. The two investigated cohorts had unbalanced groups with regard to disease status with most participants being cognitively normal and only few subjects with diagnosed AD. However, despite most participants showing no or only mild cognitive decline, especially in the ADNI cohort, we found significant associations between complexity, tau-PET and cognitive function, demonstrating that fMRI complexity is sensitive to potentially small changes in the course of AD progression. Future studies incorporating longitudinal data are warranted to estimate the rate of change from cognitive normal to AD. Furthermore, there was a significant age difference in the EEAJ cohort between the three groups. This difference could potentially bias the statistical analyses as complexity has also been shown to decline with age. To prevent such a statistical bias, age was included as a covariate in the correlation analysis between complexity and tau-PET. Most notably, the sequence protocols for fMRI data acquisition were different for the EEAJ and ADNI cohorts. While ADNI used a standard EPI with a longer TR, the EEAJ project used a modern multi-band EPI with a sub-second TR. This leads to a significant methodological difference since MSE scales for these two cohorts reflect distinct temporal frequencies. For ADNI the investigated scale represents low-frequency signal at 0.056Hz whereas for EEAJ the same scale has a frequency of 0.23Hz. Finally, because our focus was on investigating the use of fMRI complexity as an approximation of tau-PET, we did not assess the effect of Aβ on the interaction or mediation.

To conclude, our findings show that there is a statistically significant total effect between MSE and cognitive decline that is largely mediated by tau-PET, especially in temporal areas. These results support the relevance of temporal lobe tauopathy on cognitive decline ^13^. Moreover, complexity of rs-fMRI is associated with both regional tau protein accumulation and cognitive decline, and thus could provide a safe and cost-effective alternative marker for regional neuropathology and prediction of cognitive decline in AD.

## 5. Acknowledgements

Data collection and sharing for this project was funded by the Alzheimer’s Disease Neuroimaging Initiative (ADNI) (National Institutes of Health Grant U01 AG024904) and DOD ADNI (Department of Defense award number W81XWH-12-2-0012). ADNI is funded by the National Institute on Aging, the National Institute of Biomedical Imaging and Bioengineering, and through generous contributions from the following: AbbVie, Alzheimer’s Association; Alzheimer’s Drug Discovery Foundation; Araclon Biotech; BioClinica, Inc.; Biogen; Bristol-Myers Squibb Company; CereSpir, Inc.; Cogstate; Eisai Inc.; Elan Pharmaceuticals, Inc.; Eli Lilly and Company; EuroImmun; F. Hoffmann-La Roche Ltd and its affiliated company Genentech, Inc.; Fujirebio; GE Healthcare; IXICO Ltd.; Janssen Alzheimer Immunotherapy Research & Development, LLC.; Johnson & Johnson Pharmaceutical Research & Development LLC.; Lumosity; Lundbeck; Merck & Co., Inc.; Meso Scale Diagnostics, LLC.; NeuroRx Research; Neurotrack Technologies; Novartis Pharmaceuticals Corporation; Pfizer Inc.; Piramal Imaging; Servier; Takeda Pharmaceutical Company; and Transition Therapeutics. The Canadian Institutes of Health Research is providing funds to support ADNI clinical sites in Canada. Private sector contributions are facilitated by the Foundation for the National Institutes of Health (www.fnih.org). The grantee organization is the Northern California Institute for Research and Education, and the study is coordinated by the Alzheimer’s Therapeutic Research Institute at the University of Southern California. ADNI data are disseminated by the Laboratory for Neuro Imaging at the University of Southern California.

## 6. Funding

This project was funded by NIH 1R01AG066711 (Jann/Wang). Data for EEAJ cohort was acquired under NIH funds U01AG051218 and R01AG062007 (Ringman)

## 7. Competing interests

The authors report no competing interests

## Notes

### Competing Interest Statement

The authors have declared no competing interest.

